# Antifragile therapy

**DOI:** 10.1101/2020.10.08.331678

**Authors:** Jeffrey West, Bina Desai, Maximilian Strobl, Luke Pierik, Robert Vander Velde, Cole Armagost, Richard Miles, Mark Robertson-Tessi, Andriy Marusyk, Alexander R. A. Anderson

**Affiliations:** Integrated Mathematical Oncology Department, H. Lee Moffitt Cancer Center & Research Institute, Tampa, FL, USA; Department of Cancer Physiology, H. Lee Moffitt Cancer Center & Research Institute, Tampa, FL, USA; University of South Florida Cancer Biology PhD Program, Tampa, FL, USA; Wolfson Centre for Mathematical Biology, University of Oxford, Andrew Wiles Building, Oxford, OX2 6GG, UK; Department of Physics & Astronomy, University of Southern California, Los Angeles, CA, USA; Department of Mathematical Sciences, Air Force Academy, Colorado Springs, CO, USA

## Abstract

Antifragility is a recently coined word used to describe the opposite of fragility. Systems or organisms can be described as antifragile if they derive a benefit from systemic variability, volatility, randomness, or disorder. Herein, we introduce a mathematical framework to quantify the fragility or antifragility of cancer cell lines in response to treatment variability. This framework enables straightforward prediction of the optimal dose treatment schedule for a range of treatment schedules with identical cumulative dose. We apply this framework to non-small-cell lung cancer cell lines with evolved resistance to ten anti-cancer drugs. We show the utility of this antifragile framework when applied to 1) treatment resistance, and 2) collateral sensitivity of sequential monotherapies.

## Introduction

The response of living organisms to changes in environmental conditions (access to resources or abundance of hazards) may be described on a continuum scale from robust to fragile. In scarcity, fragile organisms are likely to go extinct, while only robust organisms survive. Robustness can occur at multiple scales. For example, functional redundancy describes robustness of an individual to a loss of function mutation^1^, while genotypic redundancy involves the robustness to potential variation that may evolve on the population level^2^. However, this continuum misses a key element. The fragile-robust continuum must be extended beyond robustness to a situation known as “antifragile”. These are organisms that not just tolerate, but in fact *gain* from large variability in environmental conditions.

There exist many systems which might be classified as antifragile. For example, evolution by natural selection can be viewed as an antifragile process. Through evolution by natural selection, fragile species undergo extinction, while only species which are robust to volatility will survive. The system as a whole evolves toward a more antifragile state, increasingly robust — indeed, *antifragile* — to future volatility^3^. In response to hazardous conditions, evolutionary processes may select for fast evolutionary life history strategies, increasing a species’ ability to withstand harsh conditions^4^. Robustness has long been considered a near-universal, fundamental feature of evolvable complex systems^5^.

The term “antifragile” was originally coined by market strategist Nassim Taleb to describe situations in which “some things benefit from shocks [and] thrive and grow when exposed to volatility, randomness, disorder.”^6^ He notes that “in spite of the ubiquity of the phenomenon, there is no word for the exact opposite of fragile,” and proposes naming it antifragile.” Taleb’s work was motivated by financial risk management where it is often intractable to calculate the risk of large-scale yet rare events, but it is relatively simple to predict the financial exposure should the rare event occur^7^. He, thus, proposed investment strategies which *a priori* presuppose market volatility and invest in such a way as to not only be resilient to, but to gain from, inevitable fluctuations in the market.

### Is cancer antifragile?

Akin to financial shocks, cancer treatment causes dramatic perturbations to the environmental conditions in the tumor by inducing cell death, altering the vasculature and modulating the immune landscape. Moreover, through the treatment schedule the attending clinician has direct control over the timing and magnitude of these changes. The question of how to schedule treatment for optimal results has been a long standing question in cancer research, in which breakthroughs were often achieved through an integration of experimental work with theoretical models (see refs. 8–10 for detailed reviews). The first theory for treatment scheduling was proposed by Skipper et al^11^ in the 1960s, who based on *in vitro* experiments in leukemic cells concluded that cyctoxic agents kill a constant proportion of cells. In accordance with their “log-kill” hypothesis they found that administering drug at its maximum tolerated dose (MTD) as frequently as toxicity permitted was superior to daily low-dose treatments with similar or larger total doses^12^. However, while this aggressive approach in which the tumour environment undergoes extreme oscillations has been greatly successful in leukemias and has become one of the pillars of chemotherapy schedule design^13^, it has only had limited success in solid tumours (e.g. see ref. 14), in part due to treatment resistance.

Tumors are heterogeneous populations of cells with differing drug sensitivities depending on whether cells are cycling, or not^15,16^, and depending on the presence of geno- or phenotype, or environmentally mediated drug resistance. Investigations of “metronomic therapy” have shown, for example in an *in vitro* model of colorectal cancer^17^, that resistance may be better controlled through continuous low-dose treatment (see also 8, 9, 18–20). In contrast, the more recently developed “adaptive therapy”^21–23^ advocates more irregular schedules, which are driven by the tumor’s response dynamics. Adaptive therapy has been successfully applied *in vivo* in ovarian^21^ and breast cancer^24^, and in patients in the treatment of prostate cancer^25^.

### The efficacy of uneven treatment protocols

In clinical practice, it is common to adhere to a fixed treatment protocol, where a constant dose is administered periodically (i.e. weekly; see figure 1A, purple). However, there are several notable examples where the tumor is more susceptible to a “volatile” treatment schedule, improving upon continuous therapy. A volatile (or synonymously an “uneven”) treatment schedule will have a high dose followed by a lower dose (see figure 1A, green). In some settings, it may be possible to temporarily increase the dose delivered by employing intermittent off-treatment periods. For example, one study recently demonstrated the feasibility of intermittent high dosing of tyrosine kinase inhibitors (TKI) in HER2-driven breast cancers, with concentrations of the drugs that would otherwise far exceed toxicity thresholds if administered continuously^26^. Incorporating the differential growth kinetics of drug-sensitive and drug-resistant EGFR-mutant cells into an evolutionary mathematical model, one recent study has found that intermittent high-dose pulses of erlotinib can delay the onset of resistance in EGFR-Mutant Non–Small Cell Lung Cancer (NSCLC)^27^, a result which has been confirmed in vivo^28^. Another preclinical mouse model of NSCLC indicated that weekly intermittent dosing regimens of EGFR-inhibitors (Gefitinib) showed significant inhibition of tumor load compared to daily dosing regimens with identical overall cumulative dose^29^. In humans, intermittent “pulsatile” administration of high-dose (1500 mg) erlotinib once weekly was found to be tolerable and effective after failure of lower continuous dosing in EGFR-mutant lung cancer^30^.

**Figure 1.**
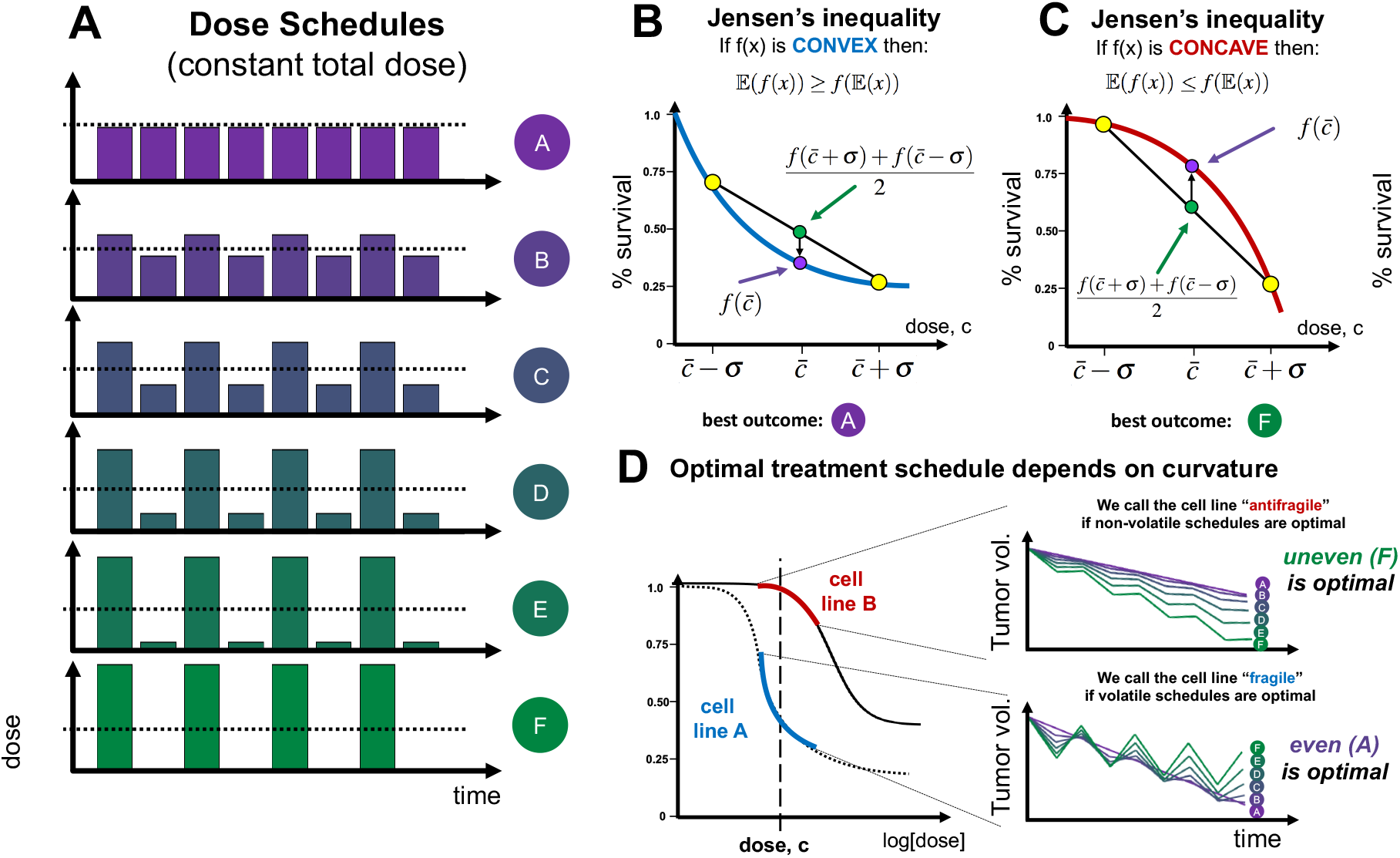
Schematic of convex and concave curvature. (A) A schematic of all dose schedules with equivalent mean dose, 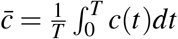 and a range of dose variance. (B,C) Example dose response curves: convex / fragile shown in B, and concave / antifragile shown in C. By Jensen’s inequality, the optimal kill is achieved by continuous, even treatment if convex (B), or uneven treatment if concave (C). (D) Optimal treatment depends on curvature. The curvature at a given dose c, may be cell line dependent (left). Curvature predicts optimal schedule (right).

In the following, we propose that the theory of antifragility provides a simple, graphical method to inform treatment scheduling. Drug response assays collect information on the response of the tumor to a range of drug doses. We will show that depending on the curvature of the drug-response relationship we can identify regions of “fragile” response in which schedules with little variability (e.g. metronomic schedules) do best, and regions of “antifragile” response in which schedules with large dose fluctuations are optimal. Subsequently, we provide theoretical as well as empirical evidence that antifragility depends on the degree of drug resistance in the tumour, and thus changes over the course of treatment. We demonstrate that as such antifragility provides a useful heuristic for informing resistance management plans, and we discuss how it can be extended to combination treatment to inform not only optimal sequencing but also scheduling of follow-up therapies.

## Methods

The purpose of this manuscript is to utilize a “fragile-antifragile” framework of dose response with the goal of determining when this continuous administration can be improved upon. The objective of this framework is to determine the treatment schedule which maximizes tumor kill. In figure 1A, we compare a range of dose schedules with identical mean dose, 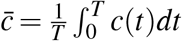 and a changing dose variance, *σ*. The continuous schedule (purple schedule, “A”) is termed an even dosing schedule, while the high / low dose schedule (green schedule, “F”) is termed an uneven dosing schedule. There are three possible scenarios to distinguish. Firstly, even schedules perform better than uneven schedules. We will call such tumors (or cell lines) “fragile.” In other words, there is no clinical benefit derived from increased treatment volatility. Conversely, it may be that uneven schedules perform better, which we will define as an “antifragile” response. Finally, both uneven and even schedules may give identical response, which we will define as a linear response. Below, we quantify the fragile and antifragile regions for a range of cell lines, and determine the benefit derived from switching to an uneven schedule in antifragile regions.

### Fragility predicts optimal treatment scheduling

In between dosing schedules “A” and “F,” there exists a range of treatment schedules (purple to green gradient; B, C, D, E in figure 1A) with identical mean dose and respective dose variance. In order to constrain each treatment schedule to an identical cumulative dose, we consider schedules which administer a pair of doses (which term a treatment “cycle”) of a high dose followed by a low dose:

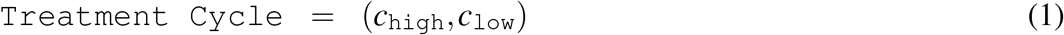

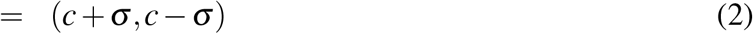

The dosing “uneveness” of a treatment schedule is given by *σ*, while the mean dose delivered is constant, *c*. Zero dosing uneveness (*σ* = 0) results in a continuous therapy of an identical dose each day. For example, we might compare an even schedule of 50mg daily to an uneven schedule (*σ* = 20mg) of 70mg followed by 30mg. The question of which is optimal is answered by directly considering the dose response curvature.

### Convexity of dose response curves

Importantly, the concept of antifragility carries a precise mathematical definition: the convexity of the payoff surface^31^. We will define antifragility (or synonymously: convexity) as follows. Let the dose response, *S*(*c*), be a twice-differentiable function of dose, *c*. The tumor’s response to treatment is antifragile if, over a range of dose *c* ∈ [*a, b*], the curvature is negative: 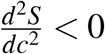. The converse implies a fragile response. For a form of this definition which more easily applicable to discrete data, we relax the assumption of differentiability and define response as antifragile if 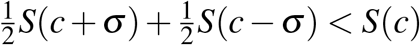. Note: the dose response indicates percent survival, and therefore a lower value is desirable (more tumor kill).

If the curve is concave (fragile) and bends downwards, this means that the value of the dose response, *S*(*c*), is greater than the average of the response of a high and low dose, 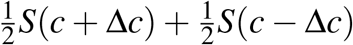. In this case, we should give the drug continuously at dose *c* (figure 1B; blue curve). Conversely, if the curve is convex and bends upwards, the value of the dose response is less than the high/low average, and the variable dosing schedule should be chosen (red, figure 1C). An explanation of this phenomena is found succinctly in Jensen’s Inequality^32,33^. If *X* is a random variable and *f* is convex over an interval in [*a, b*] then, the expected value (denoted by the symbol 𝔼) of the function is greater than or equal to the function evaluated at the expected value: 𝔼(*f* (*x*)) ≥ *f* (𝔼 (*x*)). Visually, fragility is determined by the curvature of the dose response: concave curvature bending downward is fragile while convex curvature bending upward is antifragile. In the supplementary information, we showcase the predictive power of dose response curvature for fixed treatment schedules (fig. S4) as well as intermittent, probabilistic dosing schedules (fig. S5).

The optimal schedule is dependent on the cell line as well as the dose considered. In figure 1D one cell line is fragile at dose, c, while another cell line is antifragile. The curvature predicts the optimal dose schedule for each cell line (right). Below are the results of dose response assays for a treatment-naive H3122 ALK-positive non-small cell lung cancer (NSCLC) cell line, confronted to 4 ALK TKIs, and clinically relevant chemotherapeutic agents and heat shock protein inhibitors (full panel described in Table 1). Cell lines individually resistant to a panel of four first-line therapies (Ceritinib, Alectinib, Lorlatinib, Crizotinib) as well as six additional anti-cancer agents were assayed to determine cross-sensitivity. In the next section this data, repurposed from ref. 34, are used to quantify the change in fragile and antifragile regions for 1) treatment resistance and 2) collateral sensitivity.

**Table 1.**
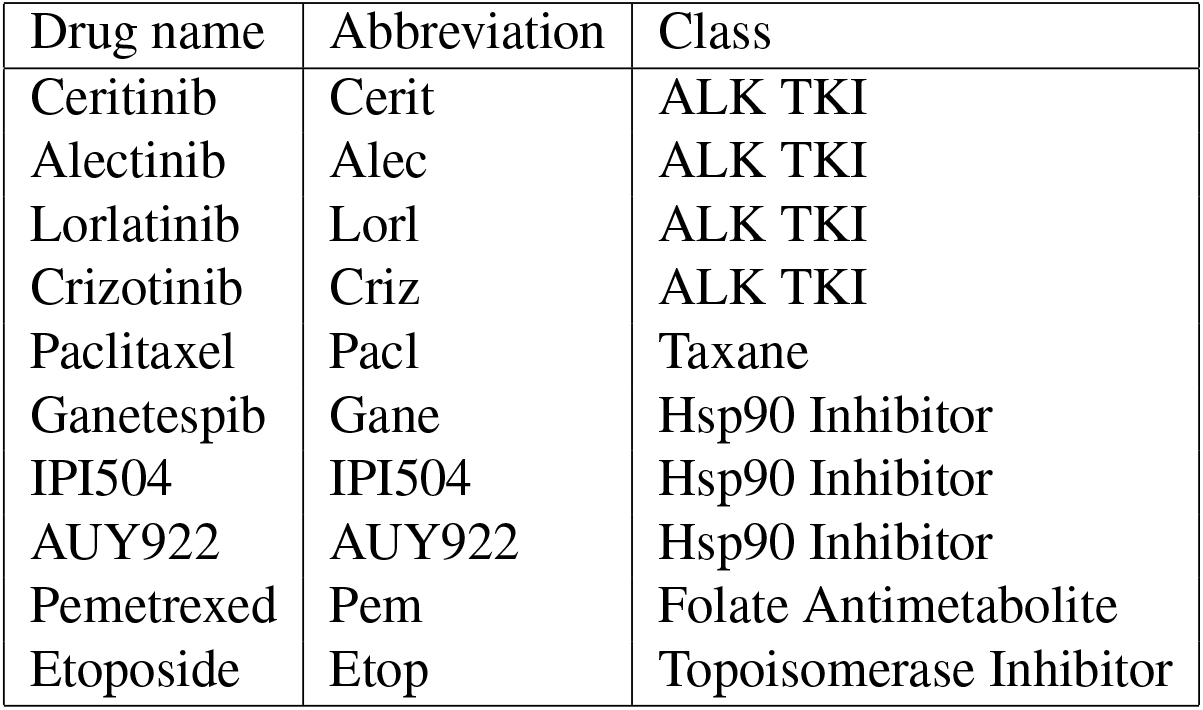

## Results

Dose response assays are often fit to the sigmoidal-shaped Hill function indicating the percent of cells which survive a given dose, *c*:

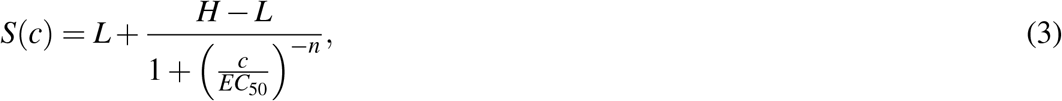

where *L* is the minimal survival proportion observed, *H* is the maximum survival proportion observed and *n* is the Hill coefficient. An example Hill-function fit is shown for treatment-naive H3122 cells confronted to Alectinib in figure 2A. Typically, a Hill function has both antifragile 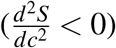 and fragile 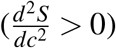 regions.

**Figure 2.**
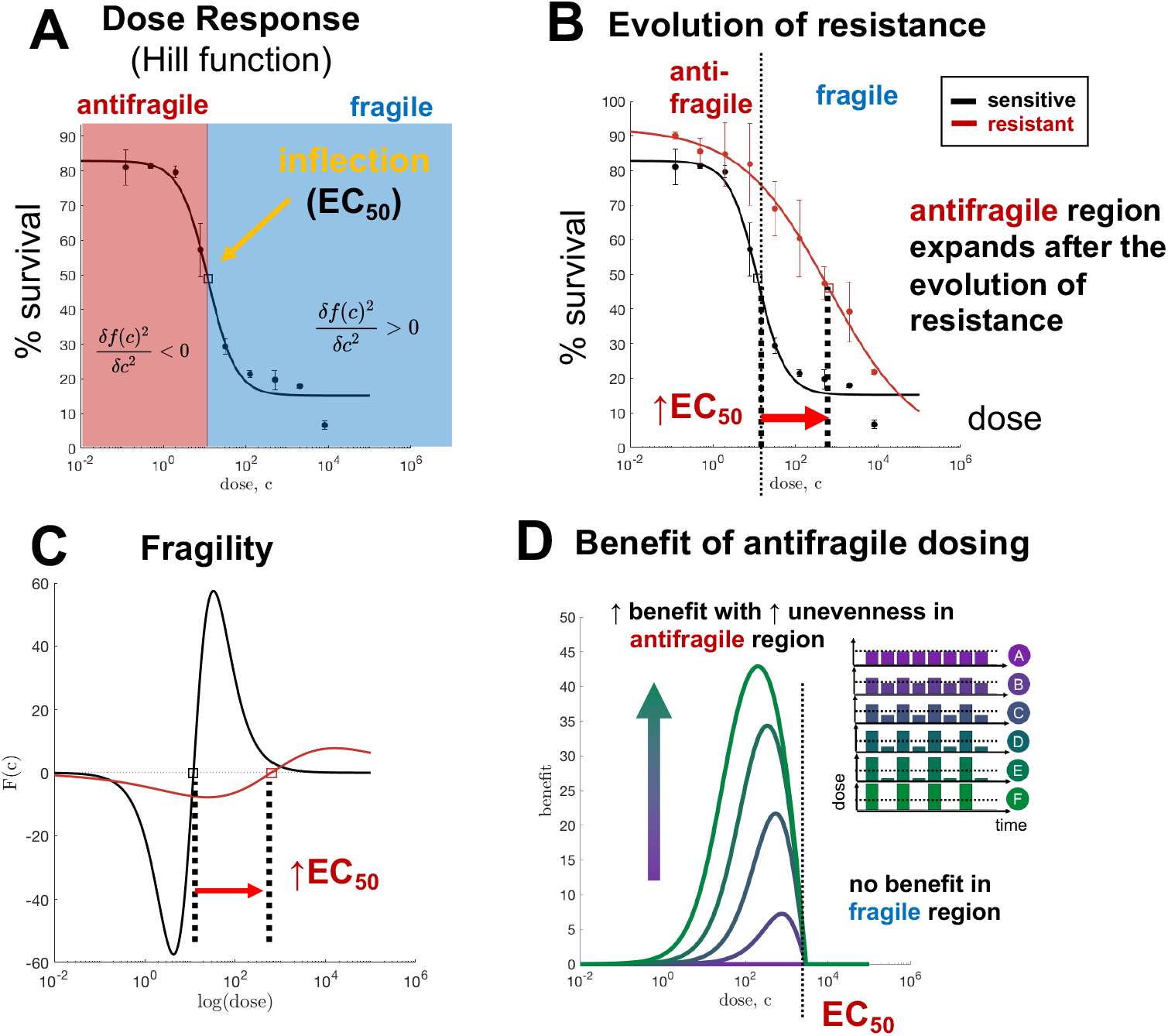
resistance. (A) Hill function best-fit (black) to HC122 treatment naive cells (circles with error bars). (B) Identical dose response assay, with evolved resistant cell line shown. (C) Fragility of naive and resistant lines, calculated from Hill function best fit. (D) Simulated percent benefit of uneven dosing schedules over continuous schedules.

### Antifragility & resistance

These treatment-naive H3122 ALK-positive cell lines were exposed to a continuous 16 weeks of drug to create a drug-resistant population, termed “evolved-resistance” cell lines. Subsequently, resistant cell lines were assayed to the same treatment. After the evolution of resistance, the dose response curve shifts from left-to-right (figure 2B, red), resulting in an increased value of EC_50_. The dose-dependent fragility of both treatment-naive and resistant cells is shown in C.

Toxicity is a limiting factor when administering treatment in cancer patients. It may not be clinically feasible to continually increase the dose administered in the manner described in eqn. 4. In figure 2 we make the assumption that higher doses are exponentially less tolerable. Mathematically, this corresponds to a treatment cycle of (10^*c*+*σ*^, 10^*c*−*σ*^), or equivalently: plotting the curvature on a log-scale x-axis. In figure 2, the EC_50_ value represents the inflection point (on a log-scale) of the Hill function. Therefore, EC_50_ is the boundary line between the fragile region (where continuous therapy is optimal) and antifragile region (where uneven treatment is optimal).

Despite their ubiquity in cancer research, dose response assays are typically used to predict and measure differential response in first-order effects, (i.e. mean value of drug dose delivered) while second-order effects (i.e. convexity) are generally ignored. In clinical practice, doses are typically administered in the fragile region (where the dose is greater than the EC50), where continuous administration is optimal. However, figure 2 clearly shows that the antifragile regions expands after the evolution of resistance, indicating that a change in treatment schedule may be necessary. In D, the benefit to switching to an uneven schedule is shown. There is no benefit in the fragile region, but significant benefit in the low-dose antifragile region. Importantly, more unevenness is increasingly beneficial (shown by the color, with dosing schedules inset).

### Fragility over time

When attempting control of a constantly evolving system, continuous monitoring and feedback is necessary to inform the timing of treatment decisions^35^. In figure 3, the rate of the loss of fragility and onset of antifragility is shown over time. H3122 cells continuous exposure to Alectinib, are assayed every week for 10 weeks. Visually, treatment-naive cell lines (week 0) exhibit a convex, fragile curvature, which flattens out as cells are exposed to treatment. Although the dose response assay is only measured for three dose concentrations here, it is still possible to calculate a discrete measure of fragility.

**Figure 3.**
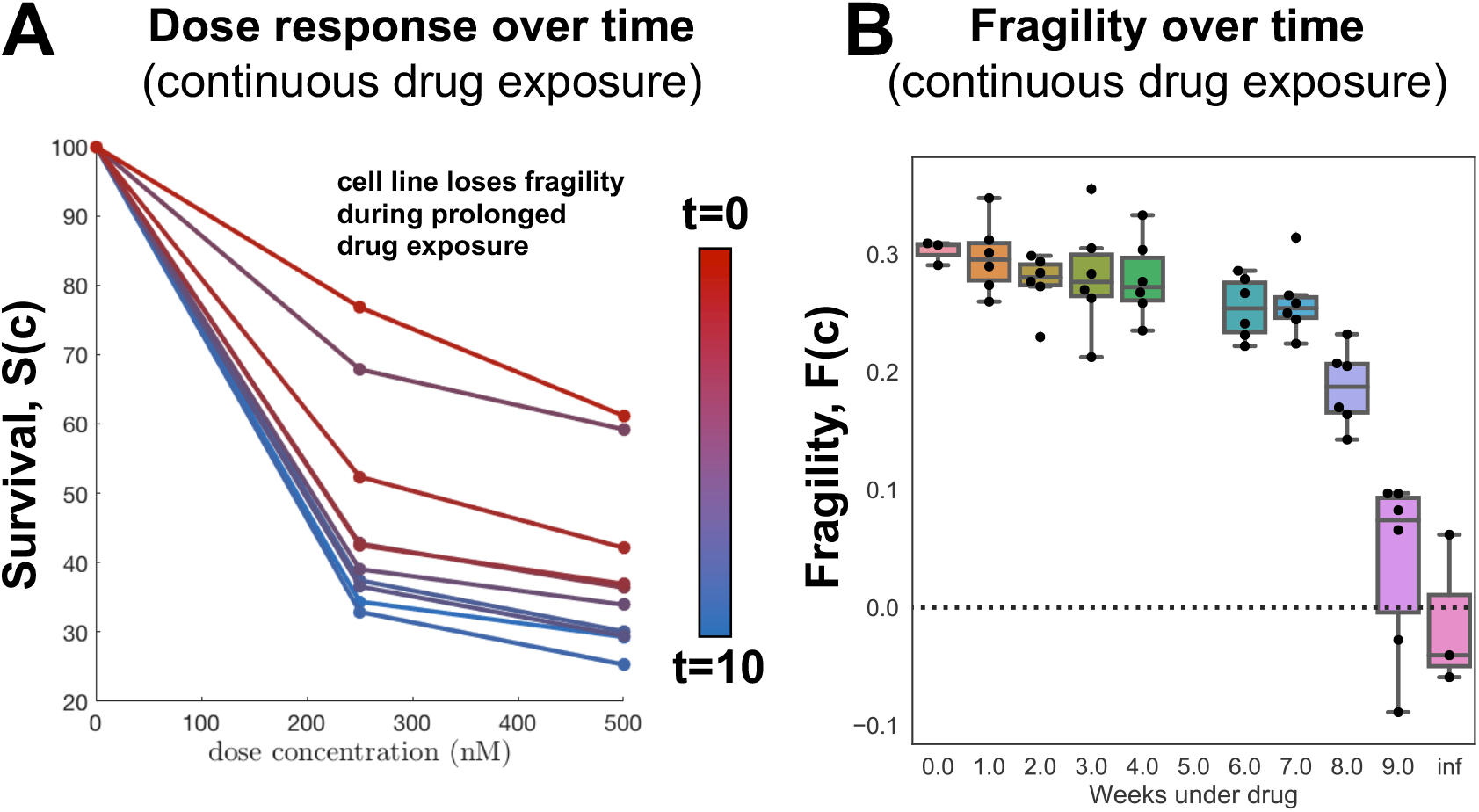
Fragility over time. (A) H3122 cells under continuous exposure to Alectinib, assayed every week for dose response at 3 concentrations (nM). (B) From A, the fragility can be directly calculated over time, with error shown for 3 biological replicates and 3 technical replicates. Note: due to an issue with data collection, week 5 is unfortunately omitted.

### Antifragility & collateral sensitivity

One proposed solution to therapy resistance may lie in finding second-line therapies which have increased drug sensitivity to the resistant population of first-line treatment. This is known as collateral sensitivity, where the resistant state causes a secondary vulnerability to a subsequent treatment which was not previously present^36^. The antifragile-fragile framework can be extended to consider the optimal dose schedule for collaterally sensitive drugs. Here, we consider monotherapy of drug 1 administered until resistance evolves, then monotherapy of drug 2. As noted previously, cell lines were cultured in continuous exposure to a range of ten treatments (see table 1) to evolve resistance. Subsequently, evolved resistance cell lines were assayed to each of the ten treatments to examine potential collateral sensitivity^34^.

Figure 4A illustrates the magnitude and sign of fragility in response to Alectinib treatment for a treatment-naive (top), as well as the full range of evolved resistance cell lines. In each row, the antifragile region is colored red, and fragile colored blue. The black line shows the boundary line of treatment naive, with arrows indicating the shift after evolved resistance to ten other anti-cancer drugs. Here, all ten treatments show an expanded region of antifragility. In other words, any of these ten treatments would enlarge the antifragile response region for Alectinib.

**Figure 4.**
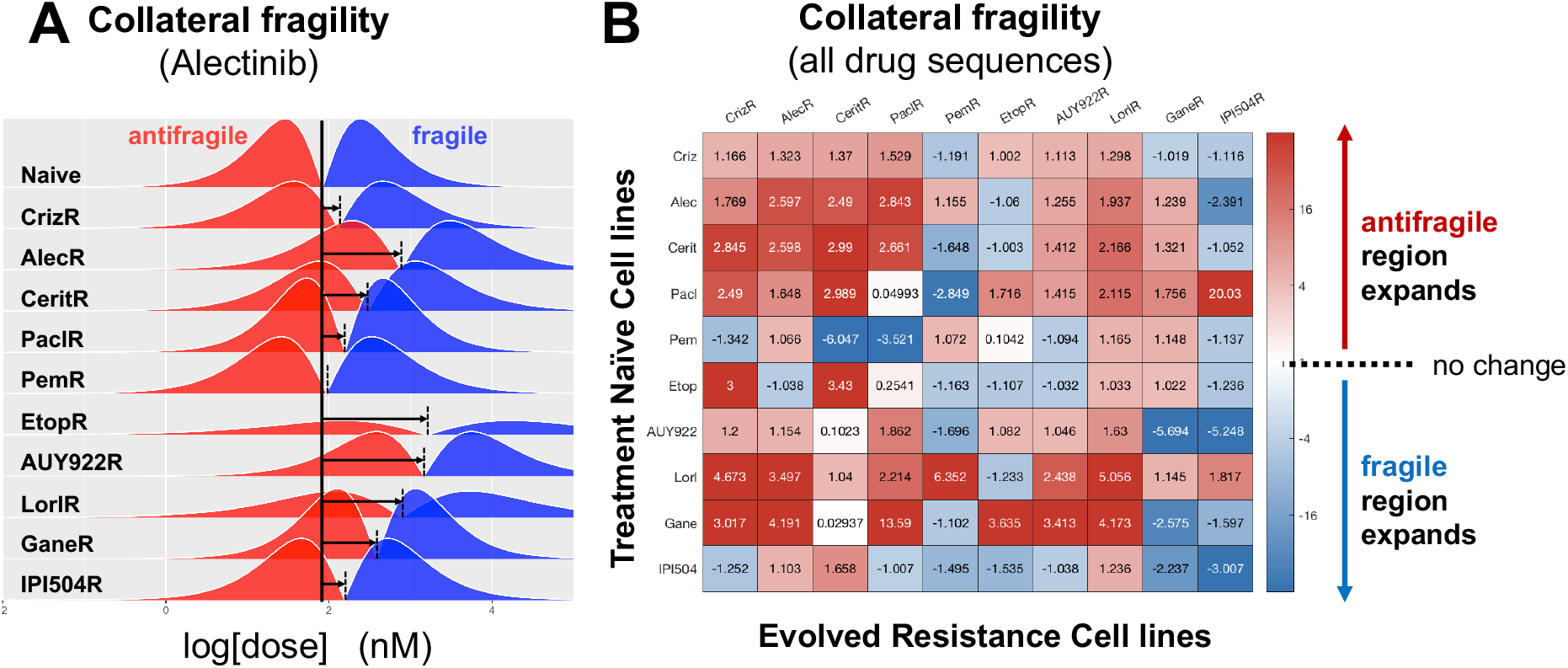
Antifragility & collateral sensitivity. (A) Magnitude and sign of dose response fragility. Vertical black line indicates the fragile-antifragile boundary for treatment-naive cells, and arrows indicate shift after evolved resistance to indicated treatment. (B) Collateral fragility: the shift for each pairwise sequence of treatments.

Likewise, every pairwise combination is shown in figure 4B, where a significant of sequential treatments are colored in red, indicating a potential improvement upon continuous therapy. The diagonal entries of this heatmap represent the shift of EC50 after evolved resistance to a single drug (as in the previous section). Seven of the ten drugs indicated an expansion of the antifragile (similar to figure 2).

## Discussion

Over the past decades it has become increasingly clear that the benefit of a cancer therapeutic agent is determined not only by its molecular action but also by its schedule. However, because of the costs associated with clinical trials and the combinatorial size of the potential search space, optimal treatment strategies remain elusive. As a result, most therapies are administered in a fashion to maximize cell kill, meaning they are given as frequently as is logistically feasible (weekly for chemotherapies, daily for orally available targeted therapies) and at the maximum dose patients can safely tolerate. At the same time, translating into the clinic alternative schedules which have been shown to perform better *in vitro, in vivo*, and/or *in silico* has been challenging, and has failed on several occasions. For example, even though “bolus-dosing” of EGFR inhibitor for EGFR-Mutant NSCLC, in which daily low dose treatment is supplemented with a weekly high dose of therapy, was shown to better control therapy resistance than the standard-of-care continuous schedule in a mathematical model^27^, as well as in *in vitro*^27^ and in *in vivo* experiments^28^, it failed to do so in patients^37^.

One reason for this discrepancy is the fact that it is often difficult to understand why a given schedule is optimal. In this paper, we have shown how that the theory of antifragility, pioneered in financial risk management, provides a general tool to compare schedules in an intuitive yet formal fashion. In particular, we have demonstrated that the curvature of the dose response curve determines whether regimens should seek to maintain a constant treatment level, or should induce fluctuations between high and low periods of exposure. Importantly, this assessment can be made graphically and does not require specialist knowledge of complex optimization techniques. Moreover, it is easily generalizable as it can be applied to dose response curves obtained from any experimental or theoretical model system.

At the same time, antifragility is supported by a thorough mathematical foundation and we have illustrated how it may be quantified in order to allow formal comparison of different cell lines or therapeutic agents. We have shown that standard-of-care for TKI inhibitors in lung cancer (continuous administration of high doses) often are applied in so-called fragile regions, affirming the optimality of standard-of-care schedules. However, our results have also shown that this conclusion breaks down after 1) the evolution of resistance, and 2) the collateral sensitivity to a previous treatment. This suggests that treatment schedules are only “optimal” for limited periods of time, and will need to be adapted as the tumour changes in response to treatment.

Treatment adjustments to manage toxicity are commonplace in clinical practice, and work on so-called “adaptive therapy” has shown that adjustments based on tumour response are clinically feasible and beneficial^25^. This treatment paradigm attempts to capitalize on competition between tumor subclones^21,38^, by maintaining drug-sensitive cells in order to suppress resistant growth due the cost to resistance. There is recent evidence that cost of resistance may be environmentally driven, depending on availability of resources^39^, and may not be required for adaptive approaches to be effective^40^. We advocate for the further development of adaptive frameworks and believe that antifragility may provide useful metric to inform when and how the schedule should be modified. Adaptive approaches typically utilize drug holidays, attempting to re-sensitize tumors during drug relaxation periods. We propose to also monitor changes in fragility after drug is removed. For example, a study adaptively administering BRAF-MEK inhibitor treatment in BRAF-mutant melanoma found that drug holidays allowed for recovery of a transcriptional state associated with a low IC-50 value^41^. This re-sensitization of drug-sensitive phenotypes during drug holidays occurs only for one cell line, WM164, while a second line showed no re-sensitization (1205Lu). Visually, the dose response curves within this study are strikingly concave (antifragile) after evolved resistance, while only the WM164’s return to a convex, fragile state after drug holiday. Other adaptive approaches utilize dose modulation, adjusting the dose higher or lower dependent on tumor response. It is still an open question on how to design effective adaptive dose modulation^42,43^. The antifragility framework introduced here may provide a path forward to predicting whether “uneven” dose modulation may outperform continuous treatment.

While experimentally validating the link between the shape of the dose response curve and treatment scheduling will be the subject of future studies, evidence for our hypothesis can already be found in the literature. Aside from the just mentioned example in melanoma, the work by Chmielecki et al^27^ provides validation in NSCLC. The authors observe a concave (antifragile) dose response curve for partially or fully resistant tumour populations treated with erlotinib. As would be predicted by our hypothesis, subsequent experiments find that bolus dosing controls resistance for longer than continuous dosing^27^.

Clinicians have a “first-mover” advantage, enabling them to exploit cancer evolution by adopting more dynamic treatment protocols which integrate eco-evolutionary dynamics into clinical decision-making^44^. While it is difficult to calculate the patient-specific risk and timing of resistance, it is often more straightforward to predict the harm incurred if a line of treatment fails. As such, treatment regimens should be designed in such a way to minimize the impact of failure, and in fact ideally turn it into an advantage. The concept that cancer evolution may be clinically steered is gaining increasing traction, and work on collateral sensitivity has already shown that resistance to one agent may induce sensitive to another^34^. In addition, we have demonstrated that along with sensitivity also the optimal mode of treatment will change. Looking ahead we propose to extend the concept of antifragility to combination therapy and to investigate, for example, the effects of drug synergism and antagonism.

To conclude, we observe that environmental fluctuations have been shown to play a role not only in treatment response but in carcinogenesis in more general. One recent consensus statement introduced the concept of an “eco-index,” a measure of hazards (e.g. drug perfusion; infiltrating lymphocytes) and resources (e.g. concentration of ATP, glucose and other nutrients; degree of hypoxia; vascular density) within the tumor ecosystem^45^. For example, instability in microenvironmental resources may lead to selection of cancer cells with fast proliferation rates^4,45^. The role of environmental fluctuations on the evolutionary dynamics of competing phenotypes has been previously studied using mathematical models of spontaneous phenotypic variations in varied nutrient conditions^46^ and of the storage effect (buffered population growth and phenotype-specific environmental response)^47^. Likewise, the Warburg effect, high glycolytic metabolism even under normoxic conditions, may arise to meet energy demands posed by stochastic tumor environments^48,49^. As such, antifragility may provide a useful metric for viewing a tumor’s response to fluctuations in environmental conditions in more general.

## Acknowledgments

The authors gratefully acknowledge funding from the Physical Sciences Oncology Network (PSON) at the National Cancer Institute, U54CA193489, as well as the Cancer Systems Biology Consortium (CSBC) grant from the National Cancer Institute (grant no. U01CA23238). Authors are also supported by the Moffitt Cancer Center of Excellence for Evolutionary Therapy.

## 1 Supplementary Information

### 1.1 Dosing schedules with identical first-order effects

As mentioned in the main text, the aim of this manuscript is to compare outcomes between even and uneven schedules. Here, a treatment “cycle” consists of two consecutive doses, 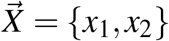. In this section, we consider schedules with identical mean cumulative dose, 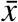, such that 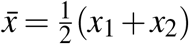. This ensures that first-order effects are constant across all treatment schedules. Let us consider treatment cycles with a high dose followed by a low dose:

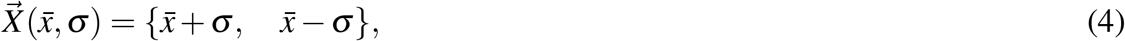

where 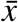 is the mean dose and *σ* is the dose variance. A value of zero for *σ* corresponds to an “even” treatment schedule.

### 1.2 Defining response

The Hill function is a commonly employed mathematical model used to parameterize dose response assays^50^. Dose response assays are experimentally determined over some time interval, *T*, (e.g. 72 hours). The Hill equation, *H*_1_(*x*), describes the survival fraction after *T* hours. We use We use *H*_1_(*x*) to describe the survival fraction for a dose, *x*:

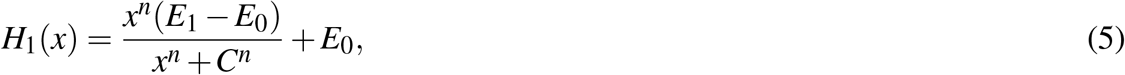

where *n* is the shape parameter (steepness of the sigmoidal curve), *E*_0_ is the response at low doses, *E*_1_ is the response at high doses, and *C* is the half-maximal response, also known as the EC50 value. Assuming cell kill is constant over the interval *t* ∈ [0, *T*], the fraction of cells killed at time *t* (where *t* < *T*) is given by:

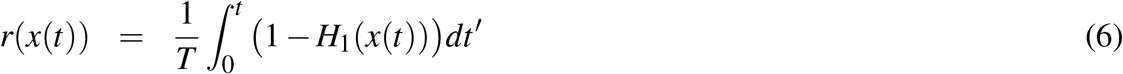

Equation 6 describes the response at time *t*, and is valid for any arbitrary dosing schedule where dose concentration is given by, *x*(*t*). Next, we consider the treatment cycle in eqn. 4, which is piece-wise constant in each half of the cycle. Total response for the full treatment cycle, 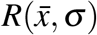, is found by adding each half treatment cycle response as follows:

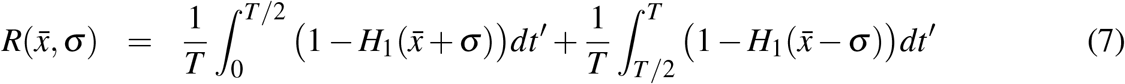

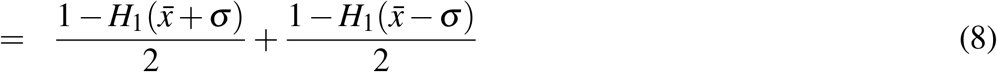

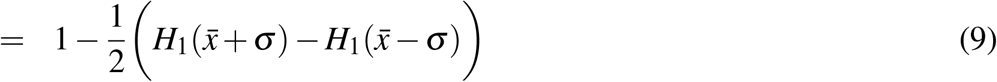

### 1.3 Defining fragility

Fragility, *F*, is defined as the gain of “uneven” high / low schedule (with corresponding unevenness parameter *σ*) over the “even” schedule where *σ* = 0:

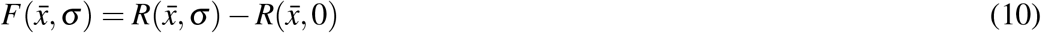

expressed in terms of the Hill function, *H*_1_(*x*), by plugging in eqn. 9:

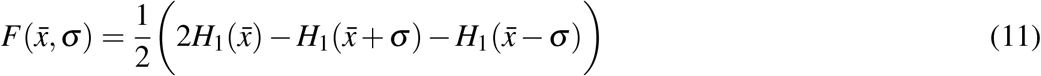

### 1.4 Fragility and Taylor Series Approximations

Next, in order to illustrate that fragilty is a “second-order” effect, we express *H*_1_(*x* ± *σ*) in terms of a Taylor Expansion about 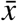:

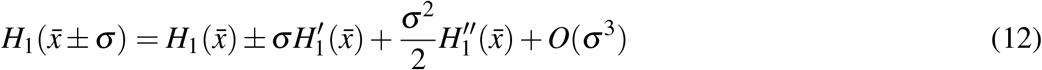

By using this expansion, fragility can be written:

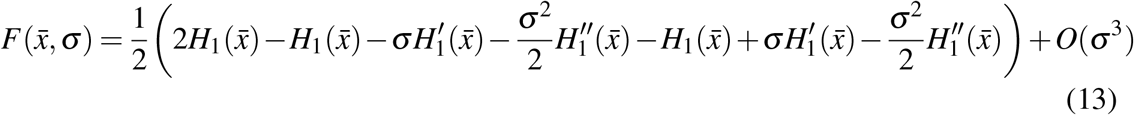

where the zeroth and first order terms cancel out to give:

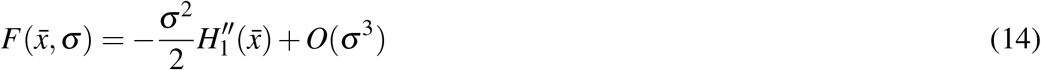

We have shown for small *σ* that fragility, *F*, is proportional to the second derivative of dose response, *H*_1_, and thus we refer to fragility as a second-order effect. In the next section, we make the connection between fragility and finite difference methods.

### 1.5 Fragility and finite differences

The derivative of a continuous, differentiable function *f* (*x*) can be approximated using finite differences. The difference quotient using central finite differences is given by:

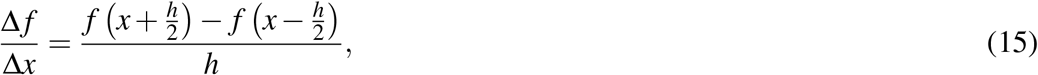

where *h* is the interval over which the finite difference is estimated (*h* > 0). This difference quotient is an approximation of the derivative *f*′ (*x*). The error of this approximation is proportional to *h*, meaning that equation 15 is a first-order approximation. By taking the limit as *h* ? 0, we arrive at the first derivative of *f* (*x*) with respect to *x*:

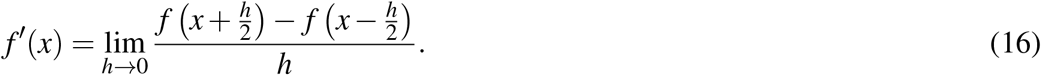

Repeating the process, we use the central difference formula for 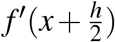 and 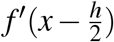 to derive the finite difference quotient for *f*′ (*x*):

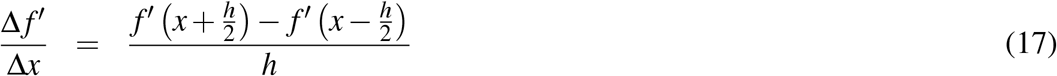

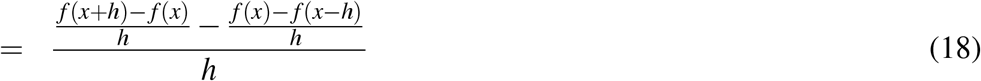

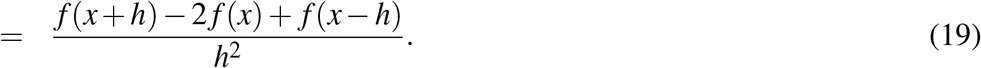

This second difference quotient (eqn. 19) is an approximation of the derivative *f″* (*x*), where the error of this approximation is proportional to *h*^2^ (e.g. see ref. 51). By taking the limit as *h* ? 0, and re-arranging terms we derive the second derivative of *f* (*x*) with respect to *x*:

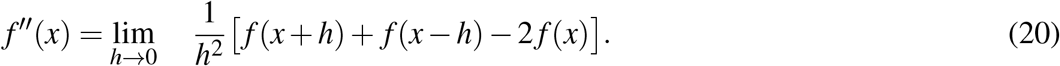

Expressing the dose response, *H*_1_(*x*), in terms of this limit gives:

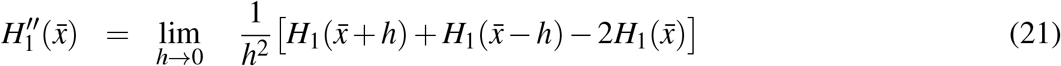

Importantly, the bracketed terms can be related to our definition of fragility, *F*, in equation 11. We obtain a relation between fragility, *F*, and the second derivative evaluated at the mean dose, 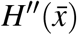:

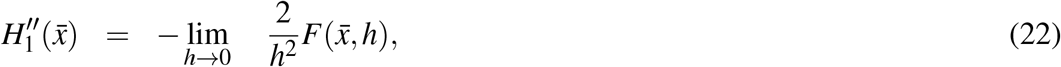

where we have substituted the bracketed term using eq. 11. Equation 22 describes the relationship between fragility and the numerical second-derivative of the function, *H*_1_(*x*). In the limit as *h* → 0, the numerical second derivative (eqn. 19) approaches 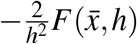. This finding indicates that the second-derivative is proportional to fragility, but only when *h* is small.

While this relationship (eqn. 22) is important to developing a foundational understanding of fragility, we are interested in comparing treatment schedules with large values for *σ*. In this case, the second-derivative is a poor approximation, so we employ equation 11. The second-derivative is accurate for predicting the gain of treatment schedules with small values for *σ*, but a poor approximation for large *σ* values.

### 1.6 Deriving the antifragile-fragile boundary

The Hill function is a commonly employed mathematical model used to parameterize dose response assays^50^. The Hill equation can be written as:

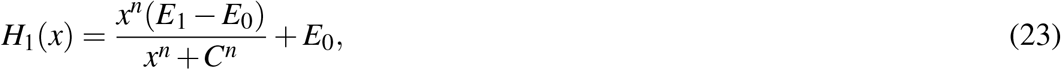

**Figure S1.**
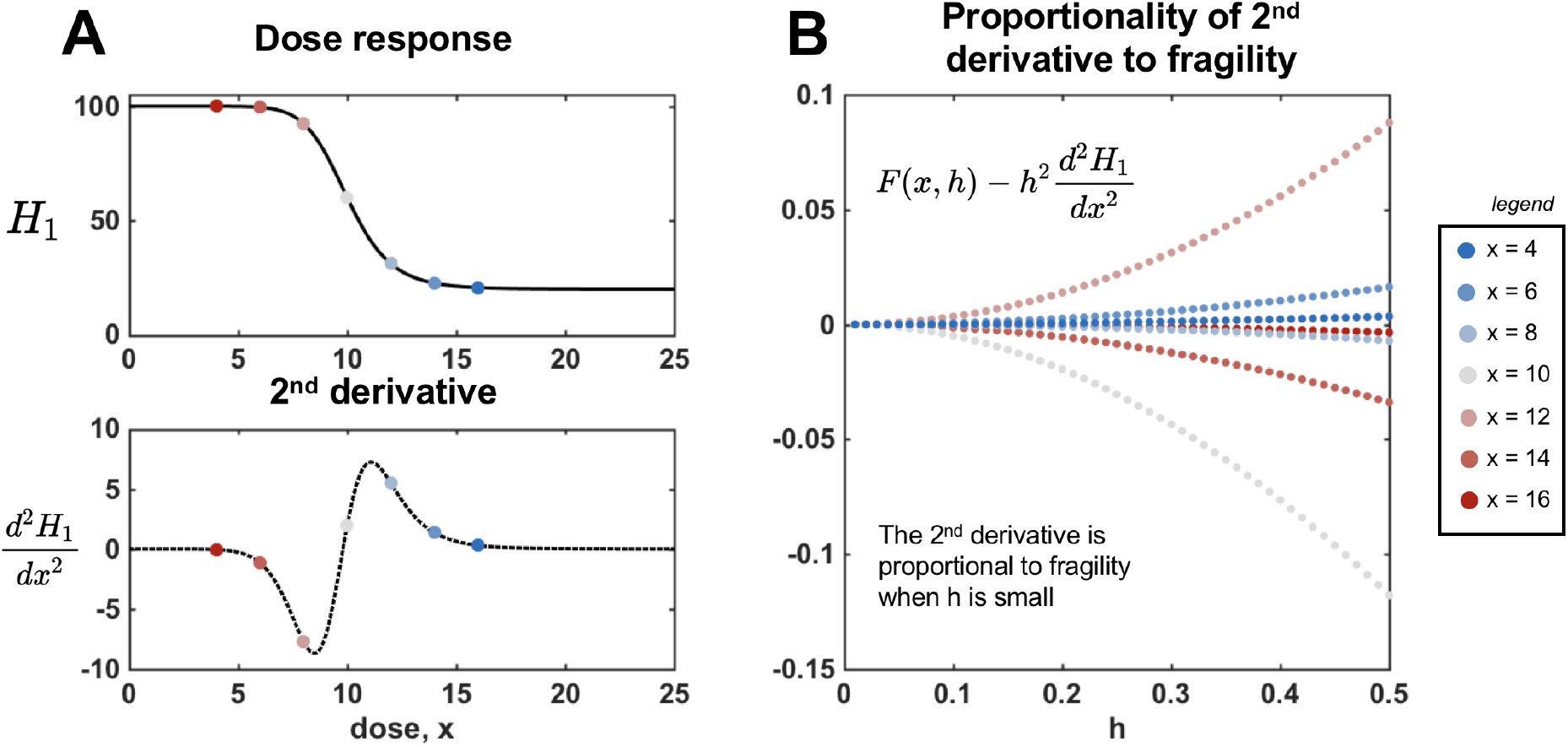
The second-derivative is an approximation for fragility for low values of h. (A) Hill function, *H*_1_(*x*) (eqn. 23) shown for *n* = 10, *E*_0_ = 100, *E*_1_ = 20, *C* = 10. Analytically derived second-derivative (eqn. 24) is shown in the bottom panel. (B) Difference between fragility and second-derivative at various dose values (red to blue) corresponding to panel A. As *h* ? 0, the error approaches zero: 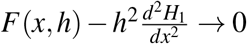.

where *n* is the shape parameter (steepness of the sigmoidal curve), *E*_0_ is the response at low doses, *E*_1_ is the response at high doses, and *C* is the half-maximal response, also known as the EC50 value. The second derivative (simplified) is:

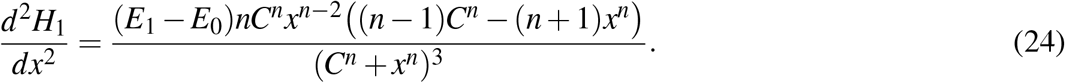

In figure S1A, an example dose response function is shown (top), *H*_1_(*x*) (eqn. 23), with corresponding second derivative (bottom). In panel B, the difference between fragility and the analytically derived second-derivative (see eqn. 24) approaches zero for small *h*, with the error scaling like *h*^2^, as predicted in equation 22.

By setting the second derivative to zero, we can solve for the dose, *x*\* corresponding to the inflection point of the Hill function. This inflection point determines the boundary between convex and concave regions.

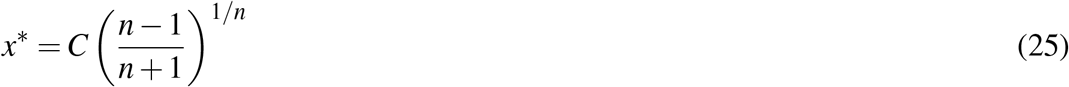

The inflection dose is a function of the EC50 value, *C*, and the steepness of the curve *n*. As seen in figure S2A, increasing the value of *n* results in moving the inflection point (blue) closer to the EC50 value (red). As *n* >> 1, the inflection point of the curve approaches the EC50 value (figure S2B). Importantly, this relationship determines the boundary between antifragile and fragile regions of the dose response curve. Benefit can thus be derived from uneven dosing if the following relation is true:

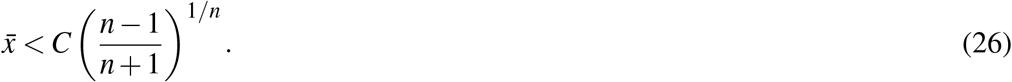

The converse implies that uneven treatment schedules provide no additional benefit.

**Figure S2.**
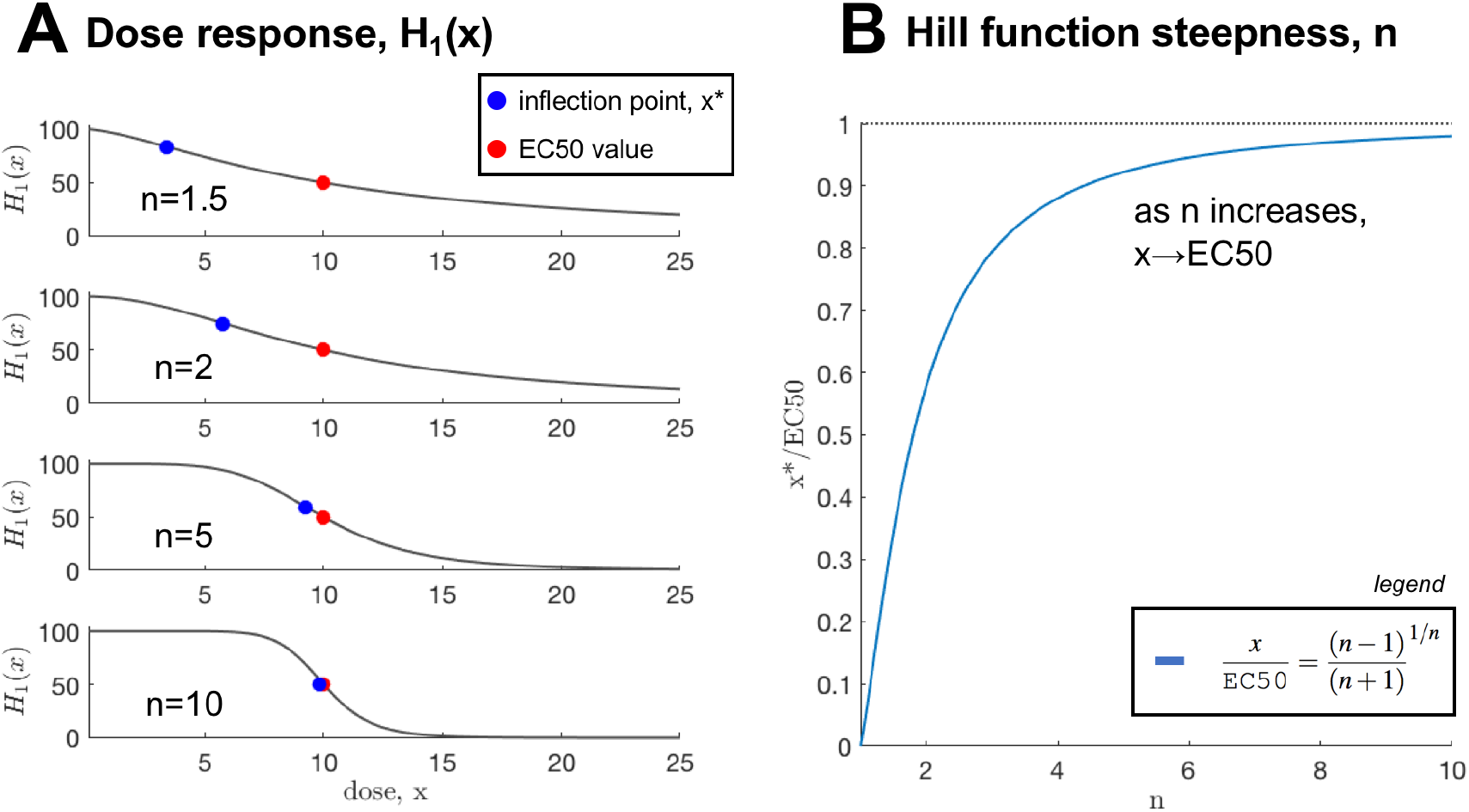
Relationship between inflection point and EC50. (A) As the steepness parameter, *n*, increases the true inflection point (*x*\*; blue) approaches the EC50 value (red). (B) The relation between inflection is derived in eqn. 25, and shown here as a function of *n*.

### 1.7 Dosing schedules with non-identical first-order effects

In the previous section, the set of treatment cycles with identical mean dose delivered was considered. In this section, we consider the fragility of a special subset of treatment schedules with non-identical mean dose delivered. Let us consider the following treatment cycle:

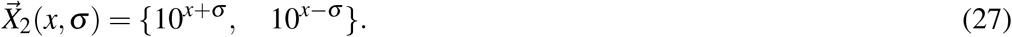

By moving each dose to the exponent, the unevenness parameter *σ* alters both the dose variance as well as the mean dose delivered: 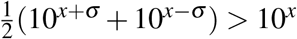. Similar to equation 11, we define the fragility of this log-transformed dosing scheme as follows:

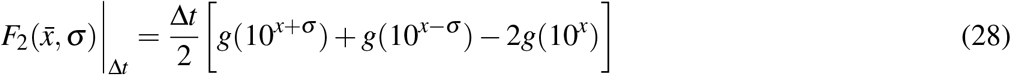

#### 1.7.1 Hill function log-transform

We refer to eqn. 30 as the “log-fragility” because the mean cumulative dose for a log-transformed variable is constant. We employ the following transformation of variables: *y* := *x*. This allows us to rewrite the Hill function as follows:

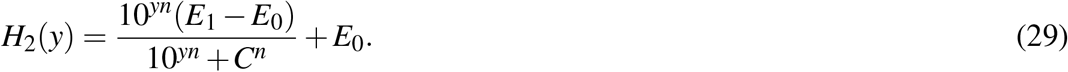

Using the transformed variable allows us to write the log-fragility as follows:

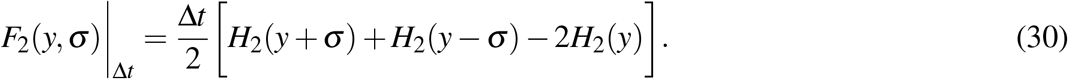

The mean of the log-transformed variable, 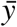 is equivalent for all values of *σ* :

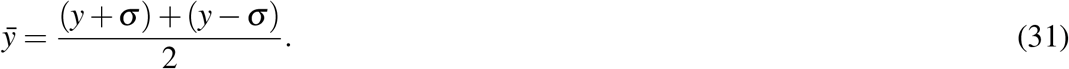

Equation 30 will provide a negative result if the Hill function is antifragile on a log-scale, and a positive result if the Hill function is fragile on a log-scale. As such, the inflection point of the Hill equation as a function of *y* will determine the antifragile-fragile boundary of our log-dosing scheme (eqn. 27). The second derivative (simplified) is:

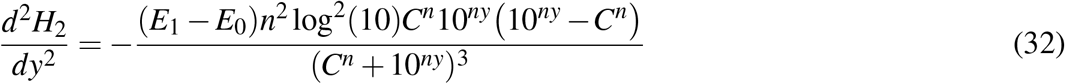

The inflection point of eqn 32 is given by:

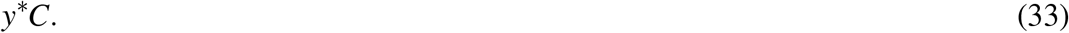

Again, this represents the boundary between the antifragile and fragile regions. Therefore, benefit can be derived from uneven dosing if the following relation is true:

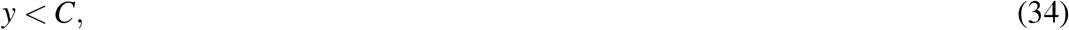

or equivalently, in our original coordinate system, benefit can be derived from uneven dosing if the following relation is true:

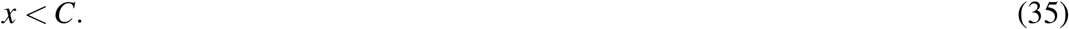

This does indeed match intuition, because the Hill function is often presented with the x-axis (dose, x) on a log-scale where the inflection point is visually represented by the EC50 value, *C*.

### 1.8 Convergence of linear and log-scale inflection

To review, we have determined the antifragile-fragile boundary for two dosing schemes: a linear system (eqn. 4) and a log-transformed dosing system (eqn. 27). Through comparison of linear fragility (eqn. 26) and logistic fragility (eqn. 35) we see that the linear case requires a lower dose for uneven strategies to be optimal. Additionally, for a large *n*, the antifragile-fragile boundary converges to *C* in both the linear and the log cases (see figure S2B).

**Figure S3.**
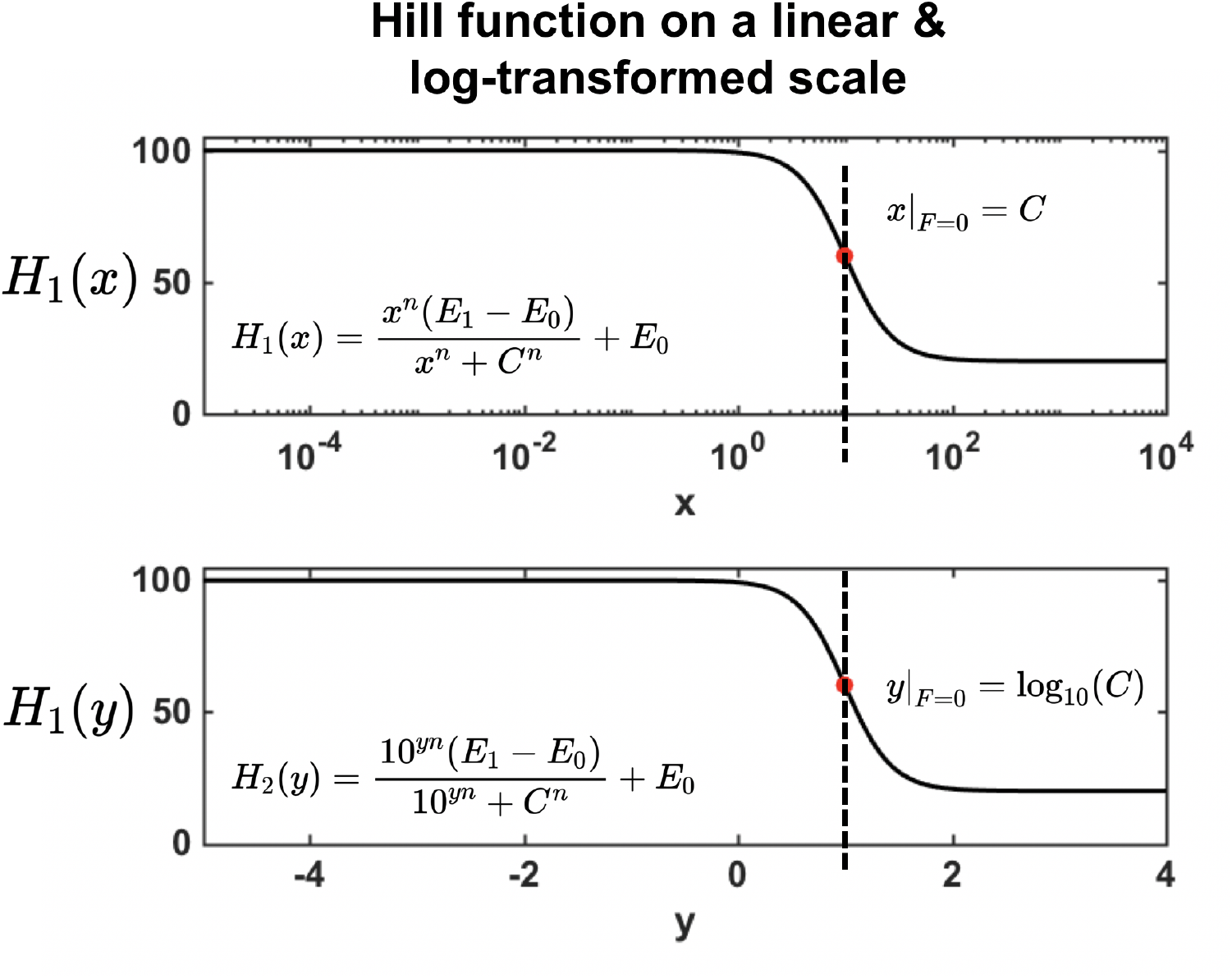
Hill function on a linear and log-transformed scale. (A) Hill function, *H*_1_(*x*) (eqn. 23) shown for *n* = 10, *E*_0_ = 100, *E*_1_ = 20, *C* = 10. The log-fragility antifragile-fragile boundary is indicated in red, where *x*|_*F*=0_ = *C*. (B) This is equivalent to displaying the Hill function in transformed coordinates (*y* := *x*) where the antifragile-fragile boundary is given by *y*|_*F*=0_ = log_10_(*C*)

### 1.9 Generalized tumor growth dynamics

To illustrate the efficacy of various fixed and random treatment dosing strategies, below we test therapeutic outcomes using a generalized model of tumor growth dynamics under treatment:

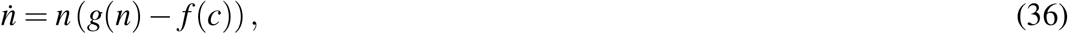

where *f* (*c*), is the fractional kill rate induced by dose *c*, and *g*(*n*) is the untreated growth rate. Here we consider exponential growth, where *g*(*n*) = *α*. Note: fractional kill, *f* (*c*), is inversely related to cell survival: *S*(*c*) = 1 − *f* (*c*). This means that the second derivative is also inverted: 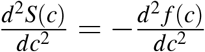. In the next section we will use this model to compare treatment schedules with identical cumulative dose.

Fixed dosing treatment schedules are simulated for two dose response functions: fragile (figure S4A; blue) and antifragile (figure S4A; red). Treatment is administered each day with varied dose unevenness (Δ*c*) but identical mean dose 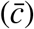. For example, in figure S4B continuous therapy (zero dose unevenness) is shown in purple, with highly uneven schedule shown in green. Tumor size over time (subject to eqn. 36) is shown in figure S4C and D. Continuous therapy is ideal for fragile dose response (figure S4C) but inferior for antifragile dose response (figure S4D).

Next, we allow the dosing unevenness to be a random variable, drawn once per treatment cycle from a Gamma distribution (probability density function shown in figure S5A), defined as follows:

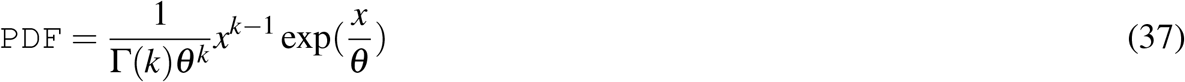

Note: the type of distribution here is not important; we choose Gamma to allow for skewed left and skewed right dosing schedules. Sample treatment schedules are shown in figure S5B, where low unevenness (approximating continuous therapy) schedules shown in purple and highly uneven in green.

The Gamma distribution is increasingly skewed right as the shape parameter, *k*, increases. Tumor size dynamics averaged over 10 treatment schedules for each value of *k*. Again, continuous therapy is ideal for fragile dose response (panel C) but inferior for antifragile dose response (panel D).

**Figure S4.**
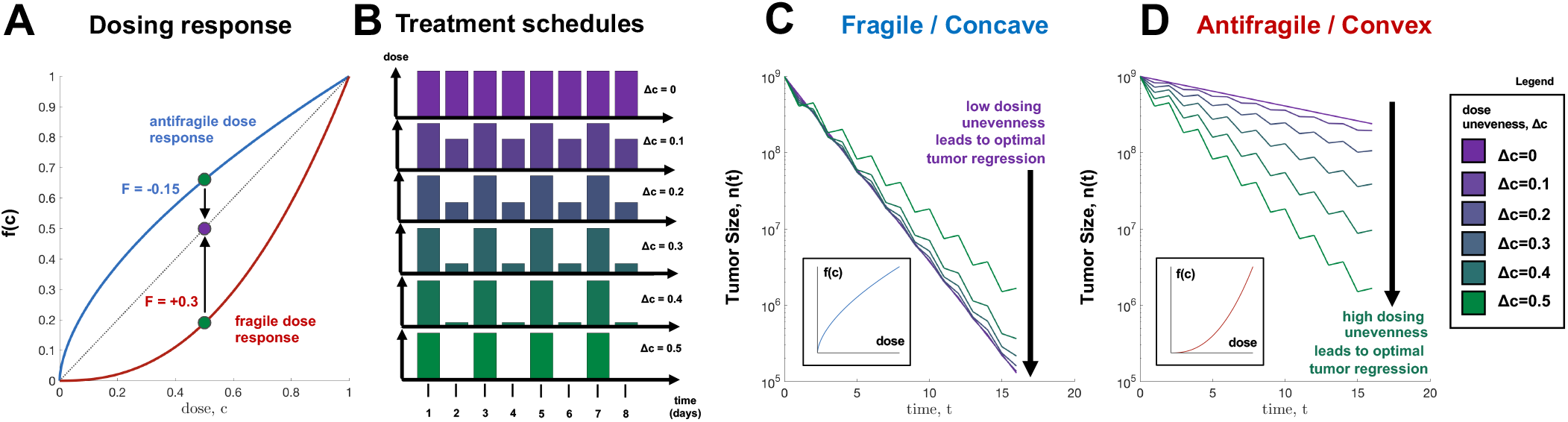
Fixed intermittent dosing strategies. (A) The dose response function (*f* (*c*) = 1 − *c*^*β*^) is fragile/concave for values of *β* < 1 (red curves), and antifragile/convex for values of *β* > 1 (blue curves). (B,C) Tumor dynamics are simulated under intermittent therapy (a dose of *c*′ + Δ*c*, followed by a dose of *c*′ − Δ*c*), colored by dose uneveness, Δ*c*. Continuous therapy (i.e. Δ*c* = 0) is shown in purple. Low uneveness (Δ*c* →0) schedules are optimal for fragile dose response curves in B while high uneveness (Δ*c* >> 0) schedules are optimal for antifragile curves in C. (D) Schematic of treatment dosing administered.

**Figure S5.**
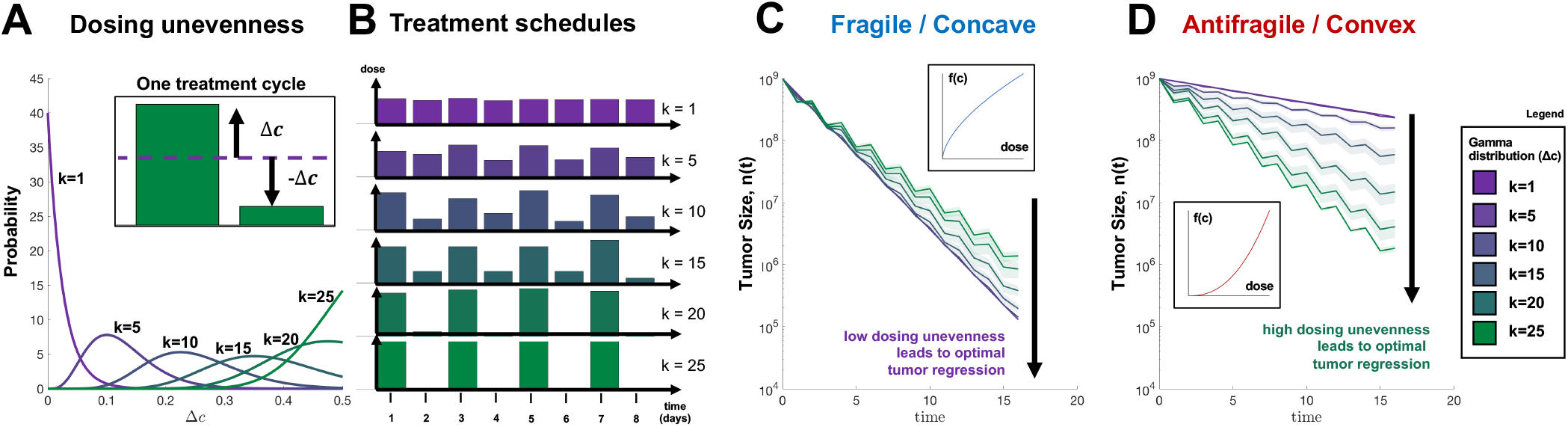
Random intermittent dosing strategies. (A) Dose uneveness, Δ*c* is now a random variable, drawn once per cycle. (B) Similar to figure S4, tumor dynamics are simulated under intermittent therapy (a dose of *c*′+ Δ*c*, followed by a dose of *c*′ = Δ*c*) for *N* = 20 cycles (averaged over 100 tumors). Continuous therapy (i.e. Δ*c* = 0) is shown in purple. Again, low unevenness (Δ*c →*0) schedules are optimal for a fragile dose response curve in (S4A) while high uneveness (Δ*c* >> 0) schedules are optimal for an antifragile curve (S4A).

